# An international scholastic network to generate LexA enhancer-trap lines for *Drosophila*

**DOI:** 10.1101/2022.11.24.517565

**Authors:** Ella S. Kim, Arjun Rajan, Kathleen Chang, Clara Gulick, Eva English, Sanath Govindarajan, Bianca Rodriguez, Orion Bloomfield, Stella Nakada, Charlotte Beard, Sarah O’Connor, Sophia Mastroianni, Emma Downey, Matthew Feigenbaum, Caitlin Tolentino, Abigail Pace, Marina Khan, Soyoun Moon, Jordan DiPrima, Amber Syed, Flora Lin, Yasmina Abukhadra, Isabella Bacon, John Beckerle, Sophia Cho, Nana Esi Donkor, Lucy Garberg, Ava Harrington, Mai Hoang, Nosa Lawani, Ayush Noori, Euwie Park, Ella Parsons, Philip Oravitan, Matthew Chen, Cristian Molina, Caleb Richmond, Adith Reddi, Jason Huang, Cooper Shugrue, Rose Coviello, Selma Unver, Matthew Indelicarto, Emir Islamovic, Rosemary McIlroy, Alana Yang, Mahdi Hamad, Elizabeth Griffin, Zara Ahmed, Asha Alla, Patricia Fitzgerald, Audrey Choi, Tanya Das, Yuchen Cheng, Joshua Yu, Tabor Roderiques, Ethan Lee, Longchao Liu, Jaekeb Harper, Jason Wang, Chris Suhr, Max Tan, Jacqueline Luque, A. Russell Tam, Emma Chen, Max Triff, Lyric Zimmermann, Eric Zhang, Jackie Wood, Kaitlin Clark, Nat Kpodonu, Antar Dey, Alexander Ecker, Maximilian Chuang, Ramón Kodi, Suzuki López, Harry Sun, Zijing Wei, Henry Stone, Anish Mudide, Chia Yu Joy Chi, Aiden Silvestri, Petra Orloff, Neha Nedumaran, Aletheia Zou, Leyla Ünver, Oscair Page, Minseo Kim, Terence Yan Tao Chan, Akili Tulloch, Andrea Hernandez, Aruli Pillai, Caitlyn Chen, Neil Chowdhury, Lina Huang, Anish Mudide, Garrett Paik, Alexandra Wingate, Lily Quinn, Chris Conybere, Luca Laiza Baumgardt, Rollo Buckley, Zara Kolberg, Ruth Pattison, Ashlyn Ahmad Shazli, Pia Ganske, Luca Sfragara, Annina Strub, Barney Collier, Hari Tamana, Dylan Ravindran, James Howden, Madeleine Stewart, Sakura Shimizu, Julia Braniff, Melanie Fong, Lucy Gutman, Danny Irvine, Sahil Malholtra, Jillian Medina, John Park, Alicia Yin, Harrison Abromavage, Breanna Barrett, Jacqueline Chen, Rachelle Cho, Mac Dilatush, Gabriel Gaw, Caitlin Gu, Jupiter Huang, Houston Kilby, Ethan Markel, Katie McClure, William Phillips, Benjamin Polaski, Amelia Roselli, Soleil Saint-Cyr, Ellie Shin, Kylan Tatum, Tai Tumpunyawat, Lucia Wetherill, Sara Ptaszynska, Maddie Zeleznik, Alexander Pesendorfer, Anna Nolan, Jeffrey Tao, Divya Sammeta, Laney Nicholson, Giao Vu Dinh, Merrin Foltz, An Vo, Maggie Ross, Andrew Tokarski, Samika Hariharan, Elaine Wang, Martha Baziuk, Ashley Tay, Yuk Hung Maximus Wong, Jax Floyd, Aileen Cui, Kieran Pierre, Nikita Coppisetti, Matthew Kutam, Dhruv Khurjekar, Anthony Gadzi, Ben Gubbay, Sophia Pedretti, Sofiya Belovich, Tiffany Yeung, Mercy Fey, Layla Shaffer, Arthur Li, Giancarlo Beritela, Kyle Huyghue, Greg Foster, Garrett Durso-Finley, Quinn Thierfelder, Holly Kiernan, Andrew Lenkowsky, Tesia Thomas, Nicole Cheng, Olivia Chao, Peter L’Etoile-Goga, Alexa King, Paris McKinley, Nicole Read, David Milberg, Leila Lin, Melinda Wong, Io Gilman, Samantha Brown, Lila Chen, Jordyn Kosai, Mark Verbinsky, Alice Belshaw-Hood, Honon Lee, Cathy Zhou, Maya Lobo, Asia Tse, Kyle Tran, Kira Lewis, Pratmesh Sonawane, Jonathan Ngo, Sophia Zuzga, Lillian Chow, Vianne Huynh, Wenyi Yang, Samantha Lim, Brandon Stites, Shannon Chang, Raenalyn Cruz-Balleza, Michaela Pelta, Stella Kujawski, Christopher Yuan, Benjamin Martinez, Reena Nuygen, Lucy Norris, Noah Nijensohn, Naomi Altman, Elise Maajid, Olivia Burkhardt, Jullian Chanda, Catherine Doscher, Alex Gopal, Aaron Good, Jonah Good, Nate Herrera, Lucas Lanting, Sophia Liem, Anila Marks, Emma McLaughlin, Audrey Lee, Collin Mohr, Emma Patton, Naima Pyarali, Claire Oczon, Daniel Richards, Nathan Good, Spencer Goss, Adeeb Khan, Reagan Madonia, Vivian Mitchell, Natasha Sun, Tarik Vranka, Diogo Garcia, Frida Arroyo, Eric Morales, Steven Camey, Giovanni Cano, Angelica Bernabe, Jennifer Arroyo, Yadira Lopez, Emily Gonzalez, Bryan Zumba, Josue Garcia, Esmeralda Vargas, Allen Trinidad, Noel Candelaria, Vanessa Valdez, Faith Campuzano, Emily Pereznegron, Jenifer Medrano, Jonathan Gutierrez, Evelyn Gutierrez, Ericka Taboada Abrego, Dayanara Gutierrez, Angelica Barnes, Eleanor Arms, Leo Mitchell, Ciara Balanzá, Jake Bradford, Harrison Detroy, Devin Ferguson, Ethel Guillermo, Anusha Manapragada, Daniella Nanula, Brigitte Serna, Khushi Singh, Emily Sramaty, Brian Wells, Matthew Wiggins, Melissa Dowling, Geraldine Schmadeke, Samantha Cafferky, Stephanie Good, Meg Reese, Miranda Fleig, Alex Gannett, Cory Cain, Melody Lee, Paul Oberto, Jennifer Rinehart, Elaine Pan, Sallie Anne Mathis, Jessica Joiner, Leslie Barr, Cory J. Evans, Alberto Baena-Lopez, Andrea Beatty, Jeanette Collette, Robert Smullen, Jeanne Suttie, Anne E. Rankin, Townley Chisholm, Cheryl Rotondo, Gareth Lewis, Victoria Turner, Lloyd Stark, Elizabeth Fox, Anjana Amirapu, Sangbin Park, Nicole Lantz, Lutz Kockel

## Abstract

Conditional gene regulation in *Drosophila* through binary expression systems like the *LexA-LexAop* system provides a superb tool for investigating gene and tissue function. To increase the availability of defined LexA enhancer trap insertions, we present molecular, genetic and tissue expression studies of 301 novel Stan-X LexA enhancer traps derived from mobilization of the index SX4 line. This includes insertions into distinct loci on the X, II and III chromosomes that were not previously associated with enhancer traps or targeted LexA constructs, an insertion into *ptc*, and eleven insertions into natural transposons. A subset of enhancer traps was expressed in CNS neurons known to produce and secrete insulin, an essential regulator of growth, development and metabolism. Fly lines described here were generated and characterized through studies by students and teachers in an international network of genetics classes at public, independent high schools, and universities serving a diversity of students, including those underrepresented in science. Thus, a unique partnership between secondary schools and university-based programs has produced and characterized novel resources in *Drosophila*, establishing instructional paradigms devoted to unscripted experimental science.

## Introduction

Conditional gene expression systems in *Drosophila* provide a powerful basis for investigating the function and regulation of genes and cells. Generation of a GAL4-based transactivator to induce expression of target genes fused to upstream activating sequences (UAS) is a widely-used binary expression system in *Drosophila* (Brand & Perrimon 1993; Hayashi et al., 2002; Gohl et al., 2011). Random genome insertion by transposons encoding GAL4 (‘enhancer trapping’; O’Kane & Gehring, 1987) generates strains with endogenous enhancer-directed GAL4 expression. Studies of many biological problems benefit from simultaneous manipulation of two or more independent cell populations or genes (reviewed in Rajan & Perrimon, 2011; Kim et al., 2021). In prior studies, parallel use of two binary expression systems allowed important new biological insights, including clonal and lineage analysis (Lai & Lee 2006; Bosch et al., 2015), ‘tissue epistasis’ studies (Yagi et al., 2010; Shim et al., 2013), and discovery of specific cell-cell interactions and contacts (Gordon & Scott 2009; Bosch et al., 2015; Macpherson et al., 2015). These approaches used a second expression system that functions independently of the UAS-Gal4 system, such as the LexA system derived from a bacterial DNA binding domain (Szüts & Bienz 2000; Lai & Lee 2006; Pfeiffer et al., 2010; Knapp et al., 2015; Gnerer et al. 2015). Fusion of the LexA DNA binding domain to a transactivator domain generates a protein that regulates expression of transgenes linked to a LexA operator-promoter (LexAop). However, the number and quality of lines expressing a LexA transactivator remains small, compared to the thousands of comparable GAL4-based lines.

To address this resource gap, we previously developed a network of partnerships between a research university (Stanford) and U.S. secondary schools to generate novel LexA-based enhancer trap drivers (Kockel et al., 2016, 2019). Here, we describe a significant expansion of this earlier effort into an international scholastic network including Stanford University, and science classes at 15 independent and public secondary schools and universities in the United States (U.S.) and United Kingdom (U.K.). This global network successfully produced hundreds of novel LexA-based enhancer trap lines for the community of science, thereby establishing science instruction paradigms rooted in experimental genetics, molecular and cell biology.

## Results

### Generation of starter fly lines for LexA enhancer trap screening

While prior studies mobilizing the X-linked SE1 element successfully generated LexA enhancer trap flies (Kockel et al., 2016, 2019), novel autosomal insertions were recovered at relatively low frequency (<5%) and showed modest expression levels of LexA. To address these limitations, we modified the SE1 element (Methods) to generate the SX4 element **(Figure 1A-B)** and SX4 ‘starter’ fly line. The SX4 P-element carries a LexA::G4 fusion (LexA DNA binding domain, “L”, the Gal4 hinge region, “H”, and the Gal4 transcriptional activation domain, “G”, construct “LHG”) identical to the SE1 P-element (Kockel et al., 2016), under the control of the hsp70 promoter. Thus, compared to the original SE1 transposon **(Figure 1A:** Kockel et al., 2016, 2019), the SX4 element has multiple modifications **(Supplemental Data File 1, Fig** 1) including (1) removal of *attB* sequences, (2) replacement of the original P-element promoter regulating LexA::G4 expression - that could lead to expression bias - with the hsp70 promoter, and (3) placement at X:19,887,268 *(amn),* a region postulated to be permissive for P-element transposition.

**Figure 1:**
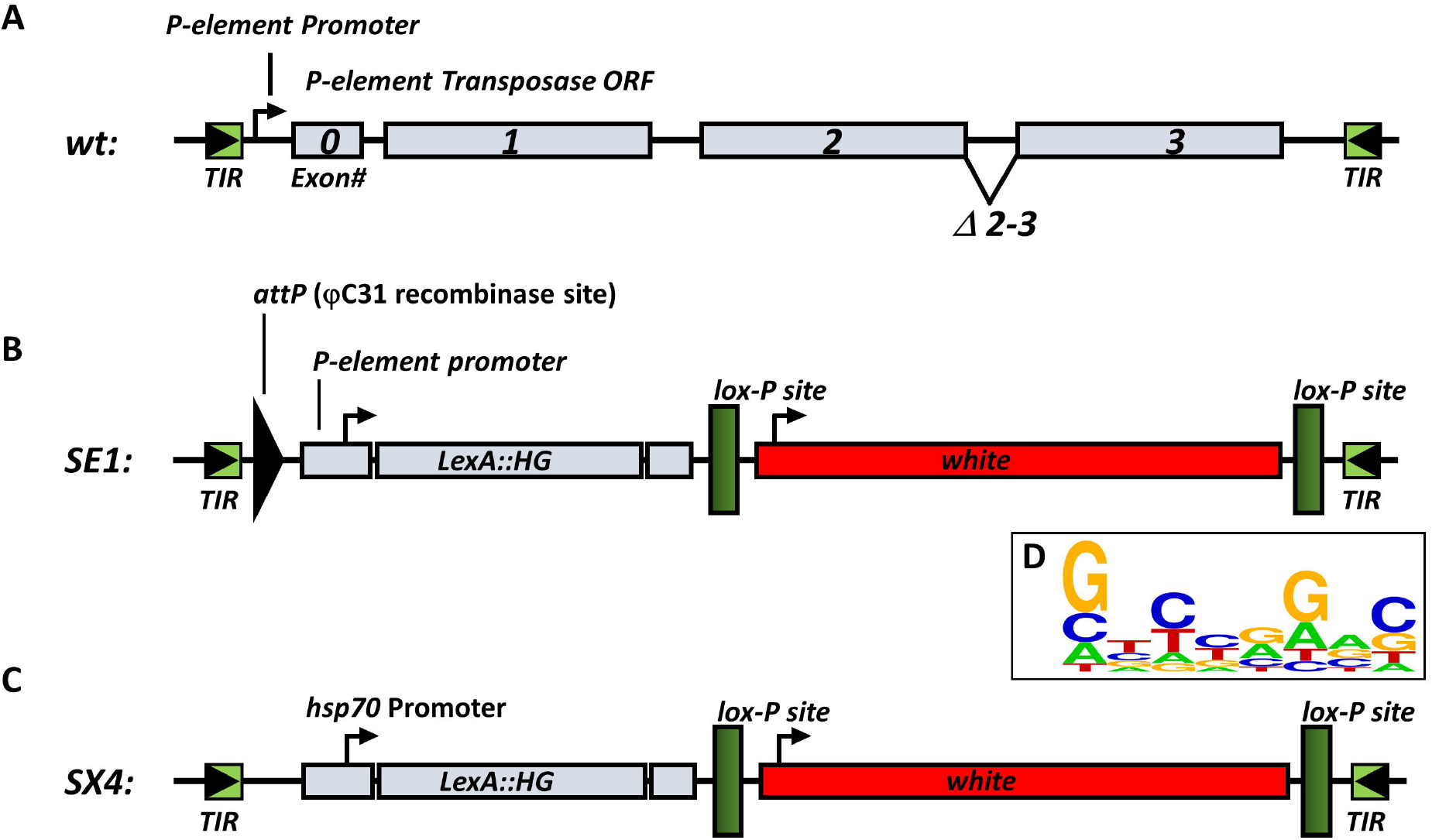
*wt* P-element and lexA enhancer traps. A) *wt* P-element described in O’Hare and Rubin, 1984. B) SE1 lexA enhancer trap used in Kockel et al., 2019. C) SX4 lexA enhancer trap used in this study. The SX4 enhancer trap encodes a lexA DNA-binding domain fused to the hinge and transactivation domain of Gal4, driven by the hsp70 promoter. The enhancer trap is marked by the *white* selectable eye color marker. See Suppl. File 1 for annotated sequence of SX4. D) Sequence Logo (see Methods) derived from 281 independent 8bp direct repeat sequences caused by SX4 insertion.

After transformation of the P{w[mC]=LHG]Stan-X[SX4]} P-element vector into the *w[1118]* strain **(Methods),** an index SX4 X-linked transformant was isogenized to the Stan-X background *iso^113232^* to generate the *w^1118^, SX4; iso#32^II^, iso#32^III^* strain. Prior studies using the SE1 element in an uncharacterized genetic background observed insertional bias of the SE1 element to a genomic region containing a KP element (Kockel et al., 2019). Thus, we used whole genome sequencing **(Methods)** with 76X and 80X coverage for males and females, respectively, confirmed the absence of KP elements in the *w^1118^, SX4; iso#32^II^, iso#32^III^* strain. Analysis of 281 8bp direct repeat sequences from individual SX4 insertions shows a slight preference of SX4 towards weak palindromic sites **(Figure 1D),** as has been reported before for SE1 (Kockel et al., 2019) and other enhancer trap P-elements (Linheiro et al., 2016).

We used the X-linked starter line to generate insertions in autosomes by hybrid dysgenesis (see below) into the isogenic background *iso^113232^*, containing the identical II and III autosomes as the sequenced strain *w^w1118^, SX4; iso#32^II^; iso#32^III^*. To complement this approach and generate enhancer trap insertions on the X-chromosome, we mobilized the SX4 element to a third chromosome balancer *(TM6B,Hu,Tb)* and generated a separate starter insertion, called *TM6B,SX4^orig^.* The *TM6B,SX4^orig^* line was used in a separate hybrid dysgenesis crossing scheme to generate X-linked enhancer trap insertions **(Methods).**

### Generating novel LexA enhancer trap lines

To generate LexA-based enhancer trap fly lines, we mobilized the X-linked SX4 P-element to isogenic autosomes *iso#32^II^; iso#32^III^* or the third chromosome SX4 insertion *TM6BSX4^orig^* to the X chromosome. To mobilize the X-linked SX4 P-element to the autosomes, we performed hybrid dysgenesis using transposase Δ2-3 at 99B (Robertson et al., 1988), to generate LexA P-element enhancer trap lines (Methods; **Suppl. Table** 1; O’Kane and Gehring 1987). Our goal was to permit interaction of the relatively weak Hsp70 promoter in the mobilized SX4 P-element with the local enhancer environment of the insertion site, and thereby generate and select LexA::HG insertions in autosomes that have spatial and temporal expression specificity (O’Kane and Gehring 1987). We used a similar experimental logic (Methods) to mobilize the SX4 located on III *(TM6B,SX4^orig^*, and isolate LexA::HG insertions in the X-chromosome **(Suppl. Table** 1).

### Characterization of Stan-X P-element insertion sites

We next used inverse PCR-based molecular methods to map the chromosomal insertion position of the Stan-X P-elements to the molecular coordinates of the genomic scaffold **(Figure 2, Suppl. Table** 1). The 301 novel insertions of this study were distributed across autosomes II and III, and their chromosomal arms (2L, 70 insertions: 2R, 69 insertions: 3L, 67 insertions: 3R, 79 insertions). We also isolated 11 insertions on the X-chromosome in a pilot hybrid dysgenesis screen using the *TM6B,SX4^orig^* as a starter line. At 19 loci, multiple P-element insertions (ranging from 2-4) mapped within 1kb, in different lines. In summary, we identified insertions at 268 unique loci, including one intergenic region **(Suppl. Table** 1); all lines were submitted to a fly stock repository (Bloomington, IN). 17 insertions were mapped to natural transposons (natural TE) present within the *iso^113232^* background. All of these natural transposons are present in multiple copies, representing repetitive DNA. 11 out of 17 *SX4* enhancer trap insertion into natural TEs could be unambiguously located to one specific site within a single copy of a natural TE. Out of these 11 natural TEs tagged by mapped *SX4, 7* are present in release6 of the *Drosophila* genome (R6, https://flybase.org/; 1360{}1206, Invader1{}757, Opus{}1033, Juan{}1190, F{}1209,mdg3{}1215, Invader4{}1371). 4 out of these 11 natural TEs tagged by mapped *SX4* enhancer trap are not represented in R6 (2x 1360, Copia, HMS Beagle, see below, **Methods).** 6 out of the total 17 SX4 insertions into natural TEs could be assigned to a specific clade of natural TEs (insertions into Doc, 2x Opus, 1731, 1360, Rt1a), but could not be unambiguously mapped within the *Drosophila* genome **(Suppl. Table** 1).

**Figure 2:**
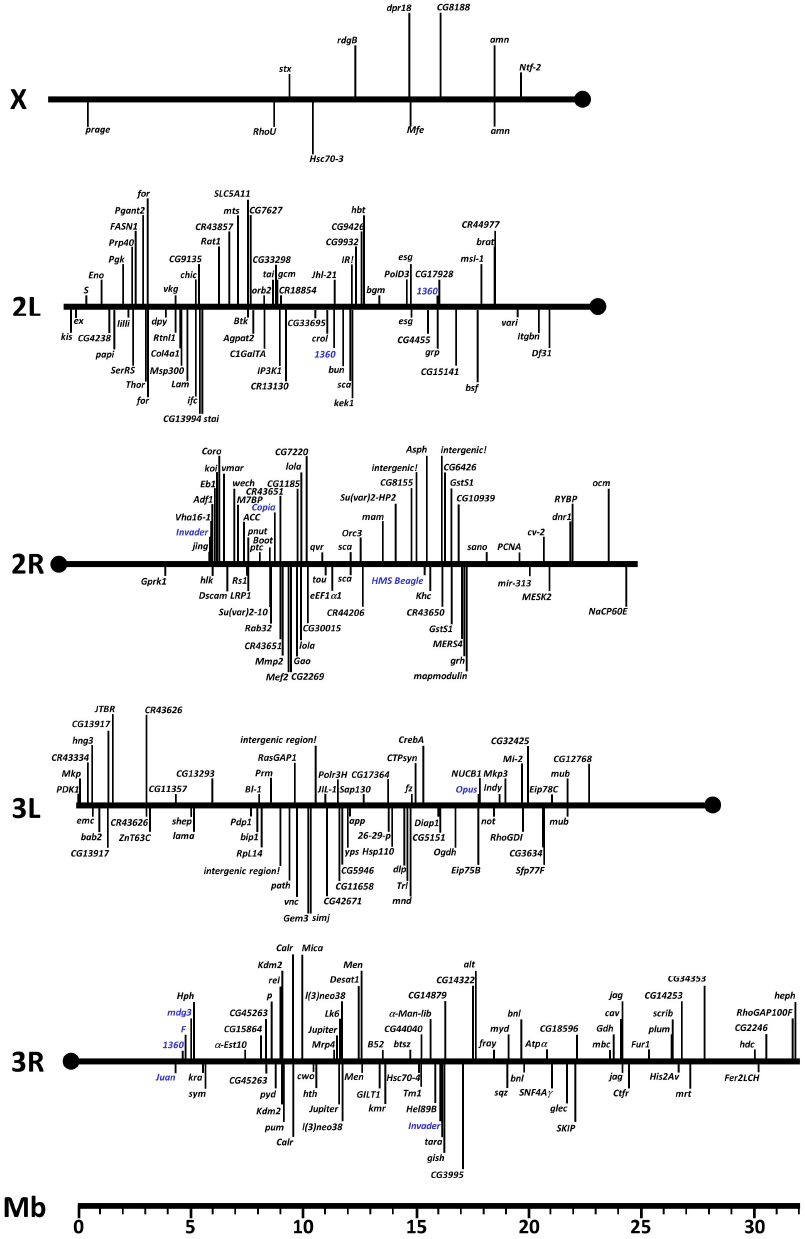
Map of novel Stan-X lexA enhancer trap insertions across chromosomal arms of X, II and III. Chromosome arms are drawn to scale and the enhancer trap positions are designated by their molecular coordinates. Scale below is in Mega-bases (Mb). P-element insertions indicated below the chromosomal scaffold are oriented 3’ to 5’, insertions above are oriented 5’ to 3’ relative to the reference sequence release 6 in FlyBase. Multiple insertions of identical orientation near identical genes are unified as single entry, and are separately listed in Suppl. Table 1. Insertions into natural transposable elements (natural TEs) are indicated in blue.

As observed previously (Bellen et al., 2011), the majority of novel insertions mapped by DNA sequencing (295/301, 98%) integrated into gene elements, including promoters, and the known first exon or intron of transcription units. Of the 295 mapped insertions presented here, we observed an even distribution of insertional direction by the SX4 P-element into genomic DNA. Using the 5’ and 3’ ends of the SX4 P-element as coordinates, we found 149/301 insertions were oriented 5’ to 3’ and 146/301 insertions were oriented 3’ to 5’. In 6 cases we were unable to determine the direction of P-element insertion. In total, we observed 17/301 (5.6%) insertions of a SX4 P-element into a natural transposon **(Suppl. Table** 1).

Natural TEs represent ~6% of sequenced euchromatin (Kapitonov and Jurka, 2003), and the percentage of SX4 insertions into natural TEs can be interpreted as a representation of that ratio. We mapped a cluster of 4 independent SX4 insertions into a variety of natural TEs *(Juan, F, 1360, mdg3:* **Suppl. Table** 1) on the pericentromeric-euchromatin boundary at 4.4 – 5.0 Mbp of chromosomal arm 3R. The small sample size precludes assignment of significance, however.

Analysis of LexA lines already present within +/-1 kb of SX4 insertions sites revealed two loci with three LexA enhancer traps previously generated, *escargot (esg)* and *SNF4γ*, which encodes the AMPK subunit gamma. These loci are known hotspots for P-element insertion, and the current study identifies three additional independent insertions into *esg*, all in the promoter region of that gene. Three SX4 insertions integrated in loci previously tagged twice with LexA insertions *(CG33298, α-Est10, lncRNA:CR43626*), and thirty-seven SX4 insertions of this study mapped within +/- 1kb of the insertion site for 1 prior LexA enhancer trap **(Suppl. Table 1).** In summary, our approach generated multiple novel LexA-based autosome and sex chromosome enhancer traps.

### Tissue expression of LexA

To verify enhancer trapping by the SX4 P-element, we intercrossed novel insertion lines with flies harboring a ‘reporter’ transgene encoding LexAop linked to a cDNA encoding a membrane-GFP (LexAop2-CD8::GFP; Pfeiffer et al., 2010), then confirmed membrane-associated GFP expression in tissues dissected from larvae **(Figure 3-5, Suppl. Table** 1). We analyzed 3rd instar larvae of bi-transgenic offspring after immuno-histochemical (IHC) staining for GFP, and simultaneous counter-staining for cell nuclei (DAPI). Image data from selected LexA enhancer trap lines were collected, and tissue expression catalogued **(Suppl. Table** 1). Within the collection, we detected GFP expression in multiple tissues of L3 larva, including neuronal cell types in the Central Nervous System (CNS), Ventral Nerve Cord (VNC) and Peripheral Nervous System (PNS), imaginal discs, and a wide range of other somatic tissues like fat body, Malpighian tubules, trachea and ring gland **(Suppl. Table** 1).

**Figure 3:**
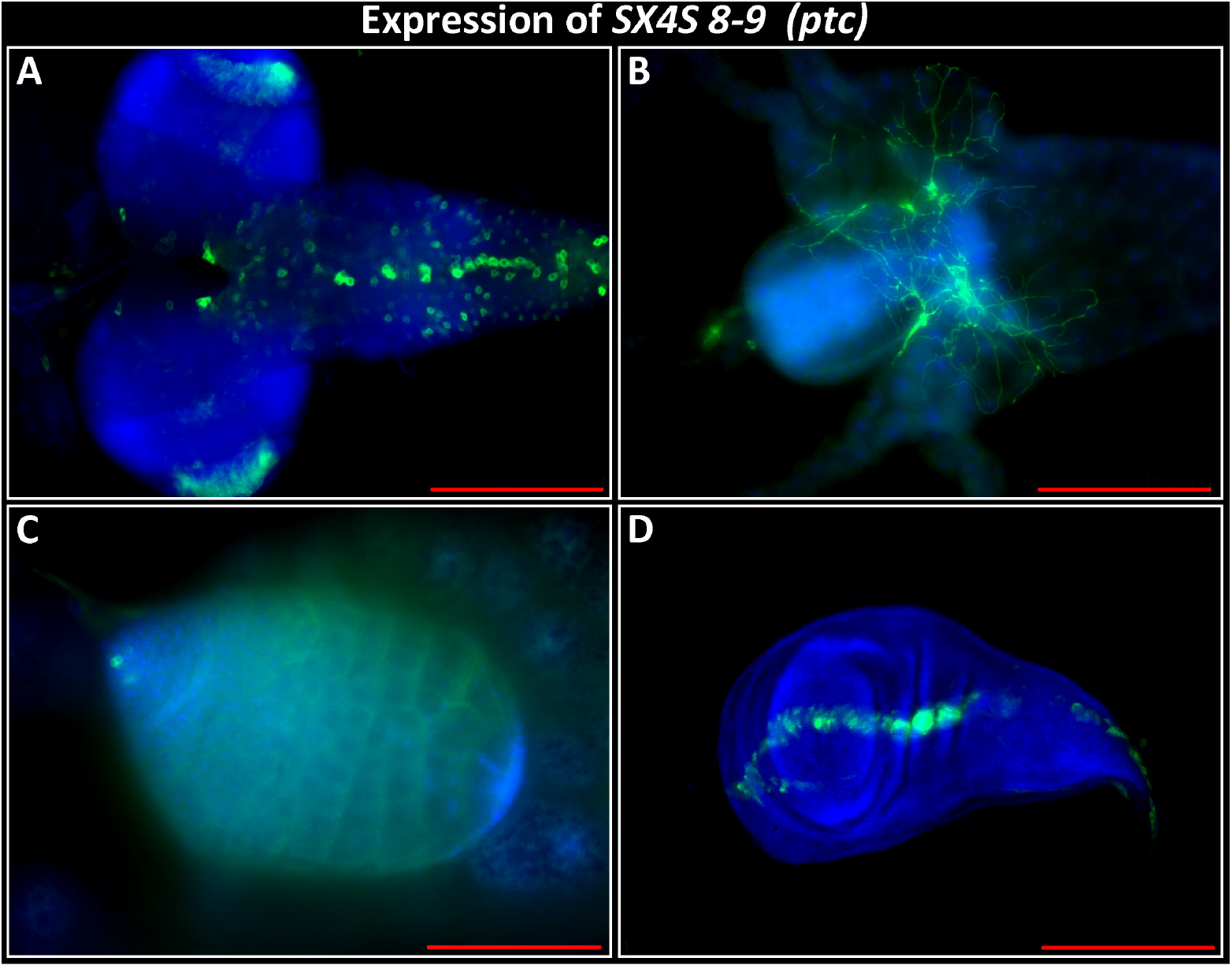
Expression of Enhancer Trap SX4S8-9, insertion in *ptc*. Genotype: *SX4S8-9/+; lexAop-CD8::GFP.* A) Third instar larval brain, expression in VNC and CNS. B) Third instar larval gut, expression in enteric neurons located at proventriculus, caeca, and midgut. C) Third instar larval testis, expression in putative hub cells and spermatocytes. D) Third larval instar wing disc. Expression along the putative anterior-posterior boundary. Anterior to the right, except D): ventral to the right. Blue: DAPI. Green: anti-GFP. Scalebar 200um, except C): 100 um

In order to test if the SX4 enhancer trap element can reproduce a known expression pattern of the gene it is inserted close to, we analyzed line *SX4S8-9*, located nearby the transcriptional start site of *patched* (ptc) at 44D1, in 2L **(Suppl Table 1, Figure 3).** Analysis of third instar wing discs revealed an expression domain along the anterior-posterior boundary, as has been described (Phillips et al., 1990). In addition, we observe expression domains in the CNS and VNC, enteric neurons, and putative hub cells and spermatocytes of the larval testis **(Figure 3,** Li et al., 2022). In summary, the SX4 enhancer trap element is able to reproduce a known expression pattern and can display activity across diverse somatic and germ line tissues and cell types.

Next, we analyzed for variety of LexA driven expression of membraned-tagged GFP in third instar larval brains from distinct SX4 insertions. For example, LexAop-CD8::GFP expression was directed by LexA from an insertion in *CG9426 (SX4Aq854), bsf/Ntf-2r (SX4Hb22-1), lola* (Sx4Lw221A) and *vnc* (*SX4Pr4*). We observed distinct patterns of cell labeling in the ring gland, CNS and VNC **(Figure 4 A-F).** To facilitate accessibility of the molecular and image data **(Suppl. Table** 1), we uploaded these to the searchable Stan-X website (stanx.squarespace.com, Kockel et al. 2016), a database searchable by expression pattern, cytology and specific genes. This includes supplementary image analysis, data from immunostaining and molecular features of SX4.

**Figure 4:**
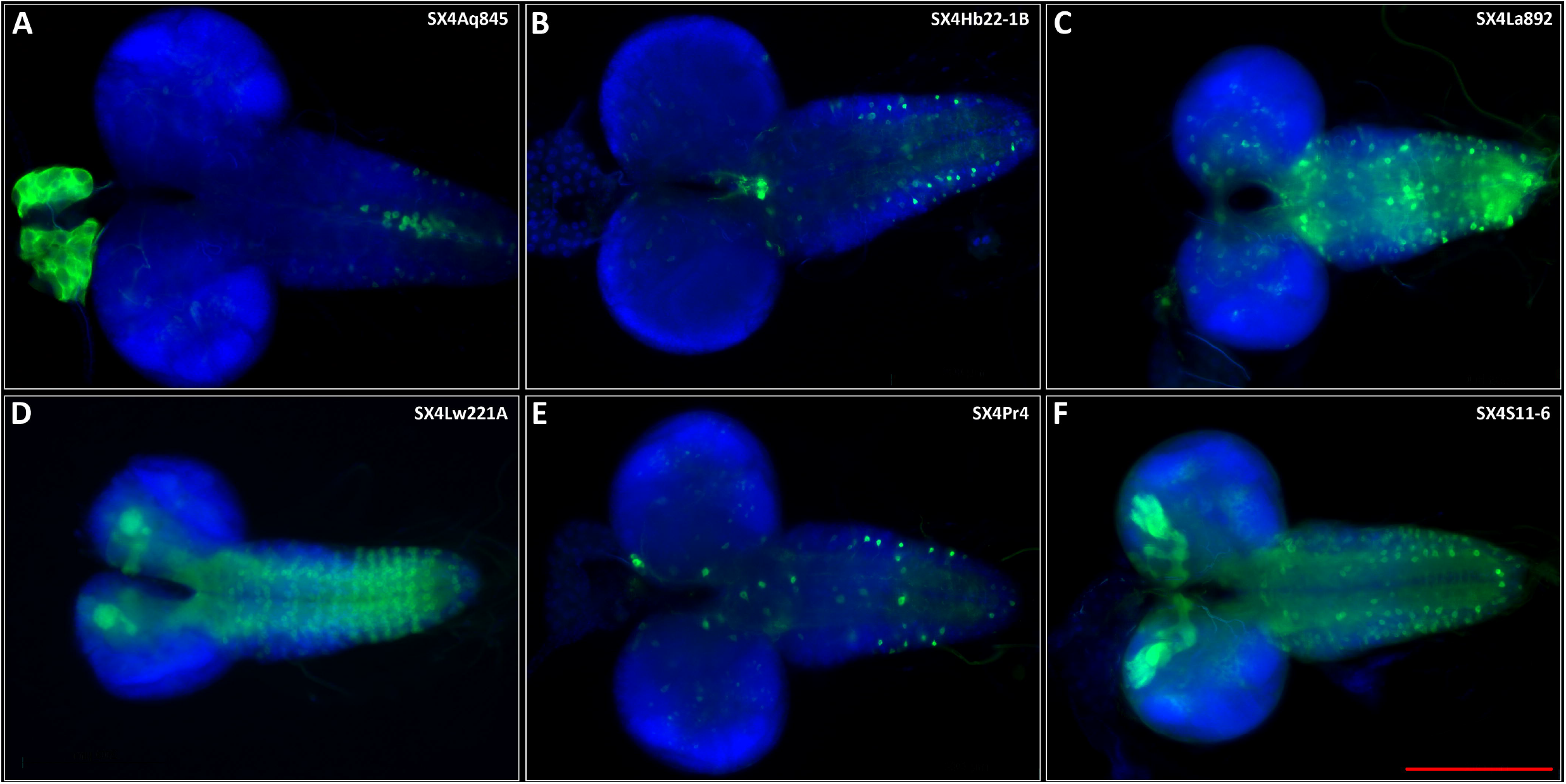
Expression pattern of 6 representative SX4 enhancer traps crossed to lexAop-CD8::GFP in wandering third instar larval brains by IHC. Green: Anti-GFP, Blue: DAPI. A) *SX4Aq845; lexAop-CD4::GFP.* B) *SX4Hb22-1B; lexAop-CD4::GFP.* C) *SX4La892; lexAop-CD4::GFP.* D) *SX4Lw221A; lexAop-CD4::GFP.* E) *SX4Pr4; lexAop-CD4::GFP.* E) *SX4S11-6; lexAop-CD4::GFP.* All images were recorded with a 20x lens, scale bar = 200um.

### Identification of SX4 lines that express in insulin-secreting neurons

Systemic insulin in *Drosophila* emanates from a paired cluster of neurons in the *pars intercerebralis* comprised of 12-14 neuroendocrine insulin-producing cells (IPCs: **Figure 5A-F).** IPCs express genes encoding *Drosophila* insulin-like peptides (Ilp’s), including *ilp-2, ilp-3* and *ilp-5* (Brogiolo et al., 2002 Rulifson et al., 2002; Li et al., 2022). Prior enhancer trap studies identified homogeneous LexA expression in these insulin^+^ cells, suggesting shared regulatory features within individual IPCs. However, we noted heterogeneous expression of the SX4Et7 enhancer trap, with expression of the LexAop::GFP reporter only in a subset of 1-2 IPCs **(green, Figure 5A)** within the cluster of Ilp2^+^ IPCs **(red, Figure 5A).** To investigate the possibility of heterogeneous genetic regulation in IPCs, we screened 87 lines and identified 16 additional lines with LexA activity in IPCs, identified by expression of the LexAop::GFP reporter in Ilp2^+^ IPCs **(Figure 5, Suppl. Table 1).**

**Figure 5:**
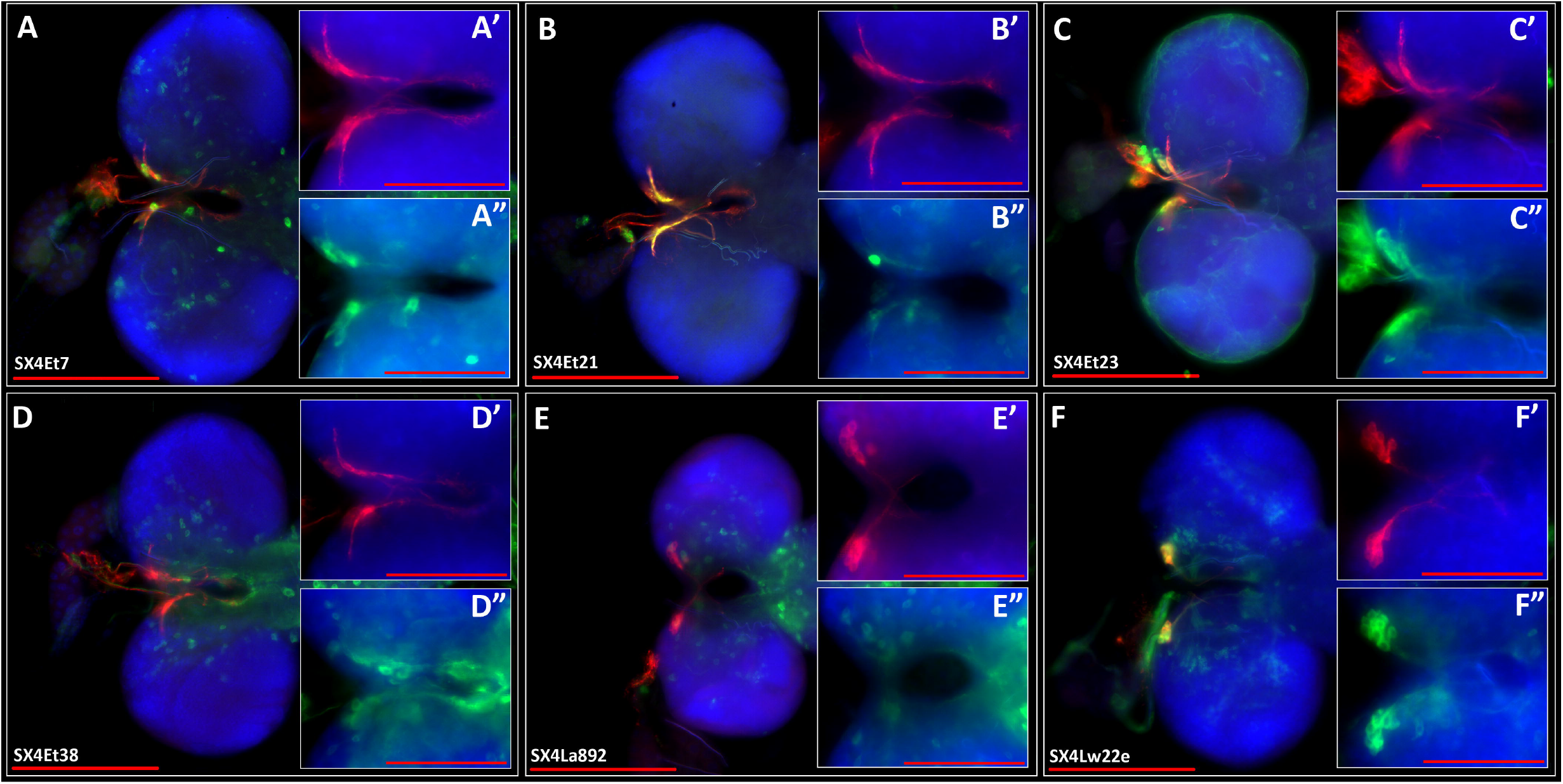
Immuno-histochemical analysis of lex-A activity in insulin expressing cells (IPCs) of selected SX4 enhancer trap lines. IPCs are marked by dilp2-Gal4, UAS-CD4::tdt (A-D) or anti-dilp2 co-stain (E-F), shown in red. Main images (A-F) were recorded with a 20x lens, scalebar = 200nm. Inserts were recorded with 40x lens (A’, A” – F’, F”), scalebar = 100nm. Green: Anti-GFP. Red: Anti-RFP (A-D) or anti-ilp2 (E-F). Blue: DAPI. A, *A’,* A”) *dilp2-Gal4, UAS-CD4:tdt; SX4Et7; lexAop-CD8::GFP.* B, B’B”) *dilp2-Gal4, UAS-CD4:tdt; SX4Et21; lexAop-CD8::GFP.* C, C’, C”) *dilp2-Gal4, UAS-CD4:tdt; SX4Et23; lexAop-CD8::GFP.* D, D’, D”) *dilp2-Gal4, UAS-CD4:tdt; SX4Et38; lexAop-CD8::GFP.* E, E’, E”) *SX4Pr5; lexAop-CD8::GFP.* F, F’, F”) *SX4Lw22e; lexAop-CD8::GFP.*

To facilitate localization of LexA activity in IPCs, we co-labeled IPCs using antibody to Ilp-2 or by specific marking of IPCs with *dilp2-Gal4* driving *UAS-CD4::tdTomato.* Of note, we observe insertions that express throughout the entire IPC cluster (e.g. **Figure 5F,** SX4Lw22e, *insertion in mayday;* **Suppl. Table** 1), and insertions that express in a subset of IPCs only (e.g. **Figure 5B,** SX4Et2, insertion in *kis).*

We selected 6 genes trapped by an SX4 insertion **(Figure 5:** SX4Et7 in *B52;* SX4Et21 in *kis;* SX4Et23 in *Afd1*; SX4Et38 in *Hel89B;* SX4Pr5 in *Star;* SX4Lw22e in *myd)* that were confirmed to drive expression in IPCs and cross-referenced expression of that gene using IPC RNAseq expression data from the Fly Cell Atlas (FCA, Li et al., 2022). The IPC FCA data is based on single-nuclei RNAseq of FACS-sorted IPC nuclei labeled by dilp2-Gal4; UAS-2xunc84::GFP (Li et al., 2022). Single-nuclei libraries from IPCs had robust expression of *ilp-2* confirming their IPC identity **(Suppl. Figure 2).** In total, data from 232 libraries from male IPC nuclei and 241 libraries from female IPC nuclei were correlated to the IHC-confirmed IPC expression data of genes tagged by the selected enhancer traps. Expression could be confirmed in all cases, in male and females, independently. The number of libraries expressing ranged from 188/232 (males) and 174/241 (females) for *kis* as the highest expressing example to 25/323 (males) and 16/241 (females) for *Adf1* as the lowest expressing example, with the number of transcripts detected correlating to the number of positive libraries. In summary, we find an overall strong correlation of genes tagged by an enhancer-trap that drive expression in IPCs and representation of RNA expression in single-nuclei RNAseq data.

### Somatic expression of natural transposons

The genome of *Drosophila melanogaster* encodes for more than 85 distinct families of natural transposons, present in multiple copies across all chromosomes (Kaminker et al., 2002, McCullers et al., 2017, **Suppl. Table** 1). Prior data suggest robust transcriptional repression of these elements in somatic and germ line cells (van den Beek, 2018, Czech et al., 2018).

We find representation of RNA derived amplicons aligning to natural TEs in the snucRNAseq libraries from IPCs and CCs (Li et al., 2022). We determine that on average ~5% and ~2% of the overall RNA content of CC cell and IPC snucRNAseq libraries, respectively, map to natural transposons **(Suppl. Figure 2).** This can be further stratified to single families of natural TEs of at least two classes, DNA cut-and-paste, and retrotransposons, for example *Invader1, Opus, Juan, 1360, F, mdg3*, and *Invader4* **(Suppl. Figure 2).** However, the presence of multiple copies of these TEs precludes the unambiguous determination of expression levels derived from a single TE element.

To address this impasse, and in order to confirm the somatic expression of natural TEs, we analyze the SX4 enhancer trap expression in L3 brains of 4 lines integrated into *1360* (two distinct elements in different locations, *SX4Ch7* and *SX4Aq839), Copia (SX4Et51),* and *HMS Beagle (SX4Et8)* natural TEs **(Figure 6, Suppl. Table** 1). These elements are specific to the *iso^113232^* background used, are not represented in R6, hence with no prior description of expression. The position and identity of the natural TE, and the insertion of the SX4 element within, was determined by iPCR **(Suppl. Table** 1) and confirmed on the genome sequence of the host genome *w^1118^, SX4; iso32^II^; iso#32^III^* by TE Mapper **(Methods, Suppl Table 2).** We find GFP positive cells in distinct patterns in brains of all four insertions **(Figure 6).** One line, *SX4Et8*, displays expression in IPCs **(Figure 6D).** Interestingly, the two independent 1360 elements tagged by SX4 insertions **(Figure 6A, B)** display distinct expression patterns, suggesting the location, rather than the TE itself, as a determinant of the pattern of expression (Treiber and Waddell, 2020). In summary, we see expression of natural TEs in snucRNASeq data of somatic cells and in enhancer trap expression of L3 brains.

**Figure 6:**
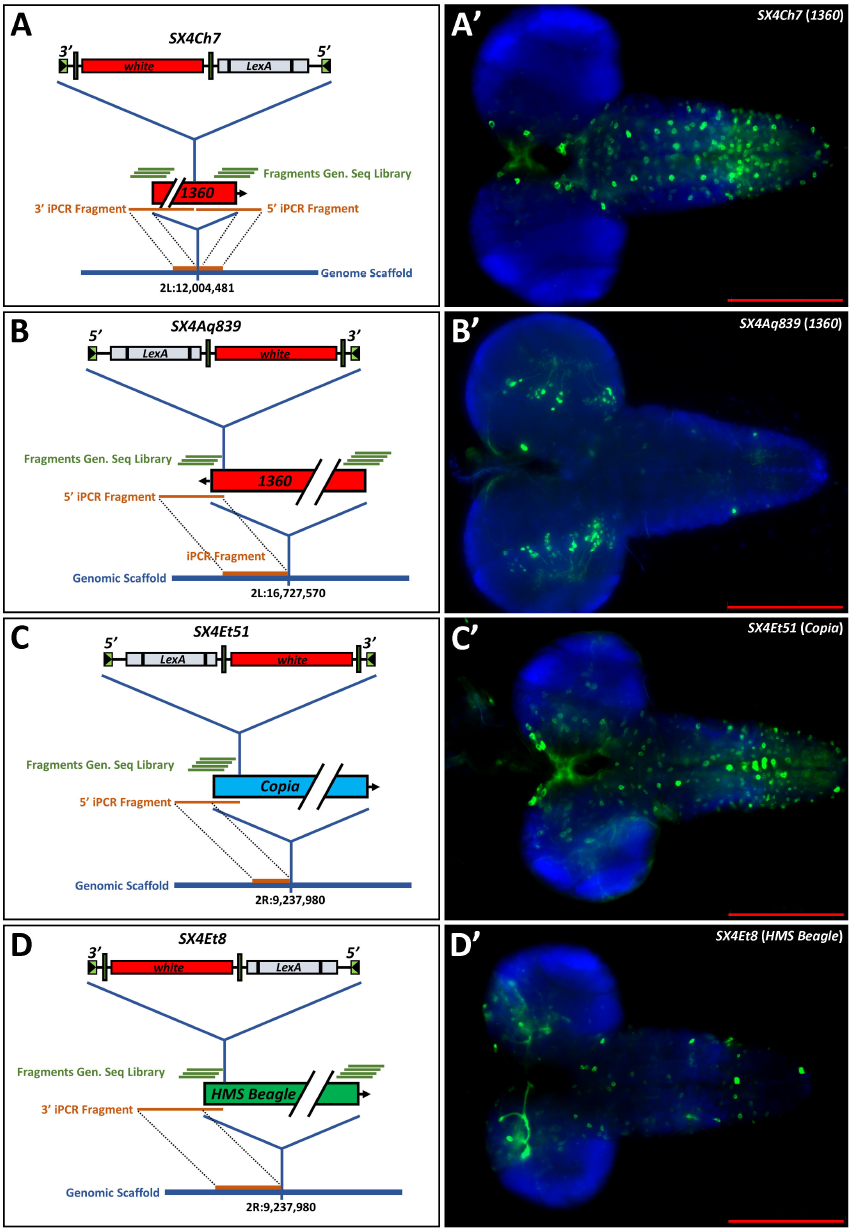
Location, tagging, breakpoint cloning and L3 brain expression pattern of natural TEs not present in FlyBase R6. A-D) Schematic representation of natural TE locations not represented in R6 and associated data. A) *1360* located at 2L:12,004,481 tagged by SX4Ch7, integrated 501 bp off the 3’ end of 1360. B) *1360* located at 2L:16,727,570 tagged by SX4Aq839, integrated 50 bp 3’ off the 3’ end of *1360,* C) *Copia* located at 2R:9,237,980 tagged by SX4Et51, integrated 174 bp into 5’ of *Copia,* D) *HMS Beagle* located at 2R:15,951,007 tagged by SX4Et8 integrated 133 bp off the 5’ end of *HMS Beagle.* Blue line: Genomic scaffold. Orange line: Sequence originated by iPCR, spanning the breakpoint of the SX4 enhancer trap and the natural TE, and the breakpoint of the natural TE with the genomic scaffold. Green lines: Sequenced amplicons from *w^1118^, SX4; iso#32^II^; iso#32^III^* genomic libraries, found by TE mapper (Methods). Sequence of amplicons are listed in **Suppl. Table 2.** Red box: natural TE. Arrow: Direction of natural TE 5’-3’. A’-D’) Third instar larval brains of respective LexA enhancer traps crossed to *w; lexAop-CD8:*:GFP. Genotypes: *A’) w; SX4Ch7/+; lexApo-CD8GFP/+.* B’) *w; SX4Aq839/+; lexApo-CD8GFP/+. C’) w; SX4Et51/+; lexApo-CD8GFP/+.* D’) *w; SX4Et8/+; lexApo-CD8GFP/+.* Blue: DAPI, Green: anti-GFP. Scalebar: 200um

**Figure 7:**
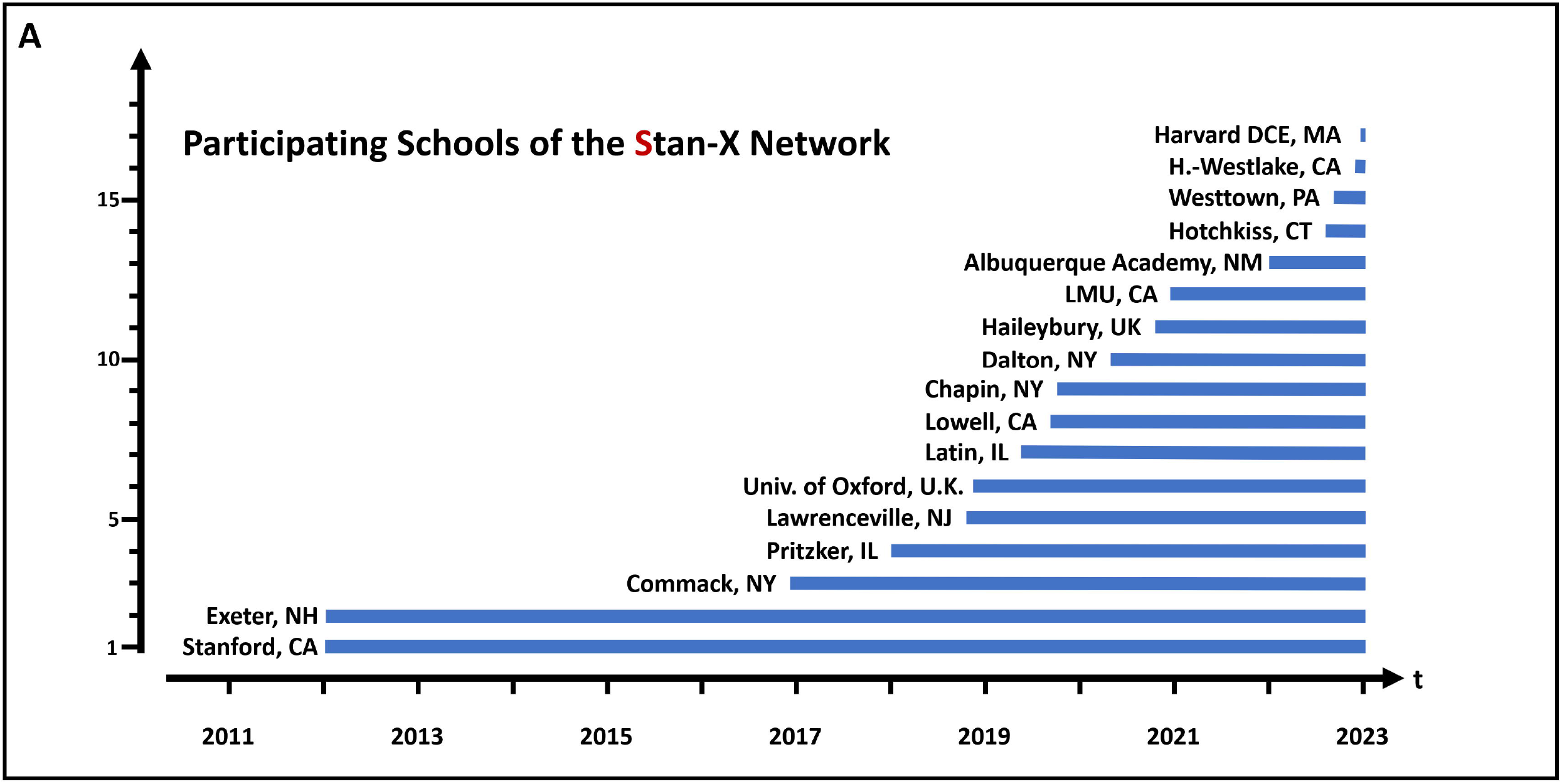
The Stan-X program. A) Timeline of school recruitment into the Stan-X program. Recruitment takes place during the school year, followed by training of the teachers in the Discover Now Teacher Academy in the summer (Suppl. Figure 1). See text for details.

### An international scholastic network to generate resources for *Drosophila* genetics

In our prior studies, we produced and characterized novel fly enhancer trap lines through an interscholastic partnership of secondary school and university-based researchers in the U.S. (Kockel et al 2016, 2019; Chang et al 2022). This involved development of curricula permitting flexible scheduling of laboratory-based ‘modules’: in some schools, this occurred in year-long courses **(Suppl. Figure 1A),** while in other schools, specific elements like mapping of P-element genomic insertions were achieved by shorter classes (Kockel et al., 2019; **Methods).** This curricular model integrated and enhanced the longitudinal quality of genetic experiments performed across years at specific schools (flies generated in one course could be characterized the following semester by another set of students). This prior work also demonstrated effective, productive collaborations of students and instructors across institutions (for example, flies generated in one school could be shared with another school that performed molecular mapping studies). Over a 10-year span (2012-2022), this group of schools collaborating with the Stanford University research group has expanded to 17 schools **(Figure 5A).**

To meet growth of the Stan-X network and demand for teacher training from 2018 on, the Teacher Academy “Discovery Now” was instituted. Incoming teachers receive a 2-week intensive training, one week online, one week in person **(Suppl. Figure 1B).** This course prepares new teachers to present the Stan-X curriculum of molecular biology and genetics to their students, and provides grounding in essential course logistics like equipment acquisition. This summer training for instructors is provided annually (www.Stan-X.org).

Here we assessed if the curriculum of *Drosophila*-based genetics, molecular biology and tissue analysis framing original, high-quality research could be adopted at additional secondary schools and universities, including abroad. As indicated by the data and resources detailed here, our studies show that research at secondary schools and universities in the U.S. and U.K. fostered production and sharing of data and fly strains, and achievement of student learning goals. In addition to curricular development at these schools, these interscholastic partners benefitted from structured interactions with network leaders at Stanford, Lawrenceville and Exeter that included weekly research teleconferences with course instructors and classes during the school year. In addition, there were university-based summer internships for students (n = 21) or instructors (n = 6), and development of annual student-led conferences with participants from multiple schools for presenting data and curricular innovations. In turn, university collaborators made regular visits to secondary school classes during the school year **(Methods).**

There were also multiple positive outcomes for students and teachers at partnering schools that were unanticipated. These included (1) emergence of *student* course alumni as *instructors* at their home institution or another Stan-X partner site (n = 6), (2) interscholastic collaboration and data development through sharing of Stan-X fly strains and other resources, (3) regular video conferencing and in-person multi-institutional student symposia organized independently by Stan-X instructors, (4) additional professional development opportunities for adult teachers, including presentation of pedagogy at professional meetings, promotion, and travel to other Stan-X partner schools, (5) development of new courses founded on CRISPR and other approaches to generate novel LexA or LexAop strains (Chang et al 2022; Wendler et al 2021) or fly genomics (see Kockel et al 2019), (6) development of Stan-X summer school courses at Harvard, Oxford, Lawrenceville and Exeter **(Suppl. Figure 1B,** https://stan-x.org), and (7) philanthropic funding for science curricular innovation and infrastructure modification to Stan-X partners. Thus, a global consortium of students and instructors at secondary schools and university-based programs have formed a unique research network actively generating novel fly strains suitable for investigations by the community of science.

## Discussion

Here, we introduced a novel lex-A enhancer trap construct in a unique isogenetic background (*iso^113232^*) and used this element to generate more than 300 novel LexA enhancer trap insertions through scholastic courses at 17 institutions. We characterized gene expression of a substantial fraction of these insertions in third-instar larval organs or tissues like CNS, VNC and gut, with a special emphasis on IPC expression in the larval CNS. We also generated a SX4 P-element on the third chromosome TM6B balancer (TM6B, SX4[orig]) that was successfully mobilized for selection of X-linked enhancer traps. In summary, these new ‘starter’ lines represent an advance over the prior SE1 LexA enhancer trap line (Kockel et al 2016).

Analysis of the insertion site sequence of these SX4 LexA enhancer traps and comparison to previous SE1 enhancer traps revealed a similar profile of insertion sequence motif. We observe a weak palindromic sequence pattern **(Figure 1D)** indicating a modest preference of Δ2-3 transposase, as observed previously (Kockel et al., 2019). Moreover, SX4 insertion directions show no preferential insertion direction, and appeared evenly distributed 5’ to *3’* and *3’* to 5’ relative to the genomic scaffold reference sequence in FlyBase, confirming earlier results. Additionally, insertion sites were enriched for regions associated with the 5’ end of transcription units, like promoters, 5’UTRs and first introns, as observed in other enhancer trap experiments (Linheiro et al., 2018). In summary, the new SX4 LexA enhancer trap element has multiple hallmarks of a P-element insertion vector.

Prior work has revealed the presence of KP elements as an impediment for hybrid dysgenesis experiments (Kockel et al, 2019). Specifically, there is a bias for precise replacement of the KP element by the enhancer trap P-element, instead of random enhancer trap insertion. This gave rise to more than 10% identical insertions stemming from replacement events with SE1 (Kockel et al, 2019). KP elements are also mobilized by Δ2-3 transposase during hybrid dysgenesis intercrosses, potentially giving rise to uncontrolled genetic heterogeneity. KP elements also encode a dominant-negative version of Δ2-3 transposase, lowering the overall transposition rate per male germ line, thereby requiring more effort to generate enhancer trap collections. Here, we used the genetic background *iso^113232^*, where we confirmed the absence of an autosomal KP element by genomic sequencing. In this background, as predicted, we observed increased transposition frequency and more random insertional distribution of our SX4 starter P-element across the chromosomes. These features have also improved workflows in participating school courses.

Here we also report the index, successful mapping of SX4 P-element insertions into natural transposons. Natural transposons are present in multiple identical, or very similar, copies per genome; thus, unambiguous mapping of insertions within these repetitive sequences requires use of modern molecular methods, including high quality iPCR and sequencing of longer DNA fragments. Specifically, these fragments need to span two ‘junctions’: the insertion site of the SX4 P-element into the natural TE, and the boundaries of the natural TE sequence with unique genomic sequence. We describe the successful mapping of 11/17 SX4 insertions in natural TE sequences, as well as the position of the SX4-tagged natural TE within the *Drosophila* genome. Of note, 4 out of the 11 natural TEs that were tagged by SX4 and positionally identified by iPCR and recipient-strain genome sequence are not represented in the current release 6 of the Drosophila genome. Therefore, they represent unique natural TEs copies specific to the *iso^113232^* background.

SX4 enhancer traps tagging natural TEs show a wide variety of somatic expression patterns in neurons of the third instar larval brain of *Drosophila*, indicating the accessibility of that locus to the transcriptional machinery of that cell. This is consistent with observations of active transcription off natural TEs in the *Drosophila* adult brain (Treiber and Waddell, 2020), the observed increase of natural TE expression in aged flies (Yang et al., 2022), and our analysis of snucRNAseq data of somatic cell types (IPCs and CC cells) from the Fly Cell Atlas. Natural TEs, in their complete form, encode their transposases (class II, cut-and-paste TEs), or other machinery mediating duplication and insertion (class I, RNA transposons, McCullers and Steiniger, 2017). In order to prevent mobilization and genome instability, the transcription off natural TEs is reported to be repressed in somatic and germ line tissues (Senti and Brennecke, 2010, Czech et al., 2018, van den Beek, 2018). However, our data suggests that somatic transcription and, putatively, transposition of natural TEs occur in somatic tissue *in vivo* (Siudeja et al., 2021, Yang et al., 2022).

Current data suggest that 6% of *Drosophila melanogaster* euchromatin is composed of natural TEs (Kapitonov and Jurka, 2003), and our natural TE insertion rate of 5.6% is consistent with this finding. We observe an insertion site cluster of 4 SX4 insertions into natural TEs at 3R, 4.4-5.0 Mbp. We also observed an insertional preference for SX4 to the 5’ and *3’* ends of natural TEs, sites of somatic transcription within natural TEs. However, our current small sample size, combined with the diversity of natural TEs, precludes the interpretation of significance of these preliminary findings.

Experimental approaches in biology benefit from temporal- or cell type-specific control of gene expression, like that possible with binary expression strategies pioneered in the Drosophila GAL4-UAS system (Brand and Perrimon 1993). Intersectional approaches, like simultaneous use of the LexA-LexAop and GAL4-UAS systems, have also greatly enhanced experimental and interpretive power in fly biology, particularly studies of neuroscience and intercellular communication (Simpson, 2016, Martín and Alcorta 2017, Dolan et al., 2017). Thus, new LexA enhancer trap lines presented here significantly expand the arsenal of available LexA expression tools (Kockel et al., 2016, Pfeiffer et al., 2013). Prior studies have demonstrated that P-element insertion in flies is non-random (O’Hare and Rubin 1983; Berg and Spradling, 1991, Bellen et al., 2011), with a strong bias for transposition to the 5’ end of genes (Spradling et al., 1995). Here and in prior work, we have found a similar preference with SX4 P-element transposition; 89% of unique insertions were located in the promoter or 5’ UTR regions of genes. Molecular characterization and studies of LexAop-regulated GFP reporter genes indicate that the enhancer traps described here are distinct, with LexA expressed in multiple tissues, including the CNS, VNC, fat body and muscle. These enhancer trap lines were submitted to the Bloomington Stock Center to enhance resource sharing.

The resources and outcomes described here significantly extend and develop the interscholastic partnership in experiment-based science pedagogy described in our prior studies (Kockel et al., 2016, 2019), which previously involved Stanford University researchers and biology classes at four U.S. secondary schools. A scholastic network, called Stan-X, now links university researchers with secondary school and undergraduate students and teachers around the world. The Stan-X network used P-element mobilization in *Drosophila melanogaster* to generate LexA enhancer trap lines reported here. Curricula based on fruit fly genetics, developmental and cell biology, and molecular biology, provided a practical framework for offering authentic research experiences for new scientists detailed previously (Kockel et al., 2016, 2019; Redfield 2012). Important research and educational goals, including a keen sense of ‘ownership’ of problems (Hatfull et al., 2006) and discovery, were achieved because the outcomes from experiments were ‘unscripted’. In addition, work permitted students and instructors to create tangible connections of their experimental outcomes (data, new fly strains) to a global science community. Data and tools from this international scholastic network demonstrated how university research laboratories can collaborate with community partners, including with resource-challenged schools serving youth under-represented in science, to innovate experiment-based STEM curricula and experiential learning that permit discovery, the *sine qua non* of science.

Indices of practical outcomes from our work include steady requests for LexA enhancer trap lines (currently >450 Stan-X lines) from the Drosophila Bloomington Stock Center, and citations noting Stan-X fly strain use in 24 publications since 2016 (e.g., Ribeiro et al., 2022; Lee et al., 2021; Zhou et al., 2020; Cohen et al., 2018). Our interscholastic partnerships and classroom-based research have expanded to include fifteen high schools and three universities on three continents **(Figure 5).** The secondary schools encompass a spectrum of public, charter, independent and ‘high needs’ schools, with day or boarding students. Three Stan-X partners are in public high schools serving ethnically and economically diverse urban communities (Lowell, San Francisco; Pritzker, Chicago; Commack, Long Island, NY), while the remainder are independent secondary schools or private universities. This experience demonstrates the feasibility and challenges of expanding the Stan-X model to public schools, which have unique resource challenges. Stan-X programs have instructed 752 students since 2012, 67% female. At independent schools, 55% of Stan-X students were female (n=562); at public high schools 70% were female (n=190). These findings suggest that curriculum-based experimental science programs like Stan-X could help address persistent gender-based disparities in science, though this possibility requires further study with case controls. Similar to the experience of others (DrosAfrica, 2020), we have found that the Stan-X curriculum can also be used abroad to foster *Drosophila-based* pedagogy abroad. Additional work outside the scope of this study is also needed to assess the longitudinal impact of programs like Stan-X on ethnic or socio-economic disparities in the scientific workforce.

In summary, this experience demonstrates the feasibility of developing productive global partnerships between schools to foster experience-based science instruction with a powerful experimental genetic organism. The thriving partnerships described here form a dynamic network of instructors, students, classes and school leaders that have produced useful science, and enhanced the personal and professional growth and development of its participants.

## Material and Methods

### Construction of the SX4 LexA enhancer-trap element

The SX4 P-element carries a LexA::G4 fusion (LexA DNA binding domain, “L”, the Gal4 hinge region, “H”, and the Gal4 transcriptional activation domain, “G”, construct “LHG”) identical to the SE1 P-element (Kockel et al., 2016), under the control of the hsp70 promoter. The 3563 bp EagI-EagI fragment from pDPPattB-LHG (Yagi et al., 2010) was subcloned to the 7097bp EagI-EagI fragment from pJFRC-MUH (Pfeiffer et al., 2010) to make pJFRC-MUH-70LHG70 (Construct #1). The 3615bp NotI-NotI fragment from pXN-attPGAL4LWL (Gohl et al., 2011) was subcloned to the NotI site on pBS2KSP vector to make pBS2KSP-attP-Pprom-GAL4-hsp70 3TJTR (Construct #2). 3842bp (NheI)-(EcoRI) fragment from pJFRC-MUH-70LHG70 (Construct #1) was Klenow filled-in and ligated to 3390bp EcoRV-EcoRV fragment from pBS2KSP-attP-Pprom-GAL4-hsp70 3’UTR (Construct#2) to generate pBS2KSP-attP-hsp70TATA-LHG-hsp70 3’UTR (Construct #3). 4098bp SacII-XbaI fragment from pBS2KSP-attP-hsp70TATA-LHG-hsp70 3’UTR (Construct #3) was subcloned to 8453bp SacI-XbaI fragment from pXN-attPGAL4LwL (Gohl et al., 2011) to generate pXN-attP-hsp70TATA-LHG-LwL (hereafter called “SX2”).

904 bp PCR product was amplified from StanEx2 using the primers XN_attP_delta_F (5’-gccgaattcggtaccGAGCGCCGGAGTATAAATAGAGGCGCTTC-3’) and LHG_R1 (5’-GCTCTGCTGACGAAGATCTACGACAATTGGTT-3’). The 1220 bp KpnI - PmeI fragment (containing the attP site) in StanEx2 was replaced by the 857 bp KpnI-PmeI fragment of the above amplified PCR product to generate pXN-hsp70TATA-LHG-LwL (hereafter, “SX4”).

The annotated primary DNA sequence of SX4 enhancer trap P-element is presented in Supplemental File 1.

### Construction of SX4 starter strains

Transformation of the P{w[mC]=LHG]Stan-X[SX4]} P-element vector into the w[1118] fly strain was performed using standard procedures. The SX4 X-linked index transformant was isogenized to the Stan-X background to generate the w^1118^, *SX4; iso#32^II^; iso#32^III^.* SX4 is located at X:19,887,269 in the *amnesiac* locus (Suppl. Table 1). We noted an Invader natural TE insertion 123 bp upstream of Stan-X[SX4] that is not represented in FlyBase rs6 of the genome, and might be specific to the *w^1118^, SX4; iso#32^II^; iso#32^III^* isogenized background used (see “Fly husbandry” below).

The SX4 X-linked insertion was then mobilized using standard procedures (see below) to the third chromosome balancer TM6B, to create the w[1118]; TM6B,SX4[orig] / ftz,e starter stain for mobilization to the X chromosome (see “Hybrid dysgenesis”). The SX4 P-element on TM6B is located at 3L: 3,250,470 in the gene encoding *lncRNA:CR43626.*

### Immuno-histochemistry (IHC)

All tissues were fixed in 4% Formaldehyde/PBS for 30 min, permeabelized in 0.2% Triton X-100/PBS for 4 hours and blocked in 3% BSA/PBS for 1 hour. All antibody stainings were performed in 3% BSA/PBS, incubation of primary and secondary antibodies were O/N. PBS was used for all rinses and washes (3x each for primary and secondary antibody incubation steps). Antibodies used: Chicken anti-RFP 1:2000 (Rockland, 600-901-379). Goat anti-GFP 1:3000 (Rockland 600-101-215). Donkey anti-Goat ALexA488 (Life Technologies, A11055). Donkey anti-Chicken Cy3 (Jackson ImmunoResearch 703-165-155). Donkey anti-Mouse ALexA594 (Life Technologies A21203). All secondary antibodies were used at 1:500. All samples were mounted in SlowFade Gold mounting medium with DAPI (Life Technologies, S36938).

### Epifluorescent Microscopy

Microscopy was performed on a Zeiss AxioImager M2 with Zeiss filter sets 49 (DAPI) and 38HE (ALexA488) using the extended focus function. Used compound Epifluorescent microscopes for high schools with all required lenses, installation services, and optional training sessions are available for sale from MicoOptics (https://www.micro-optics.com/).

### Fly husbandry and isogenized fly strains

All fly strains were maintained on a standard cornmeal-molasses diet (http://flystocks.bio.indiana.edu/Fly_Work/media-recipes/molassesfood.htm). The following strains were used: *y[1],w[1118]* (Bloomington 6598), *w[*]; ry[506] Sb[1] P{ry[+t7.2]=Delta2-3}99B/TM6B, Tb[1]* (Bloomington 1798), crossed to the Stan-X isogenic background *(iso#11[X]; iso#32[II]; iso#32[III]),* resulting in *w[1118] iso#11[X]; iso#32[II]; ry[506] Sb[1] P{ry[+t7.2]=Delta2-3}99B/TM6B,Hu,Tb[1]*, and the balancer strain *w[1118] iso#11[X]; L[*]/CyO; ftz[*],e[*]/TM6,Hu,Tb[1].* The SX4 element was first established as the X-liked index insertion of the SX4 enhancer trap P-element in a standard white background *w[1118],* producing *w[1118],P{w[mC]=LHG]Stan-X[SX4].* Subsequently, autosomes II and III of the stain were isogenized to *w[1118], P{w[mC]=LHG]Stan-X[SX4]; iso#32[II], iso#32[III].* The *TM6B,SX4[orig],Hu[1],Tb* chromosome was generated by transposition of SX4 to *TM6B,Hu[1],Tb* as described above.

### Hybrid dysgenesis from X to autosomes II and III

**F_0_**: Females of donor stock *w[1118], SX4; iso#32[II], iso#32[III]* were mated to males *w[1118] iso#11[X]; iso#32[II]; ry[506] Sb[1] P{ry[+t7.2]=Delta2-3}99B/TM6B,Tb[1],Hu.*
**F_1_**: *w[1118], SX4; iso#32[II]; ry[506] Sb[1] P{ry[+t7.2]=Delta2-3}99B/ iso#32[III]* males were crossed to *w[1118] iso#11[X]; L[*]/CyO; ftz[*] e[*]/TM6,Tb,Hu* females.
**F_2_:** *w+* males were mated to *w[1118] iso#11[X]; L[*]/CyO; ftz[*] e[*]/TM6,Tb,Hu.*
**F_3_:** The insertion line was stably balanced deploying a brother-sister cross of *w+* animals that contained *CyO* and *TM6B,Hu[1],Tb*, yielding *w[1118] iso#11[X]; CyO/SX4[#] iso#32[II]; TM6B,Hu[1],Tb/ iso#32[III]* for insertions on chromosome II, or *w[1118] iso#11[X]; CyO/ iso#32[II]; TM6B,Hu[1],Tb/SX4[#] iso#32[III]* for insertions on chromosome III.

### Hybrid dysgenesis from autosome III to X chromosome

**F_0_:** Females of donor stock *w[1118]; TM6B,SX4[orig],Hu[1],Tb/ftz[*] e[*]* were mated to males *y[1] w[1118]; CyO, PBac{w[+mC]=Delta2-3.Exel}2/amos[Tft]* (Bloomington# 8201).
**F_1_**: *w[1118]; CyO,PBac{w[+mC]=Delta2-3.Exel}2/+; TM6B,SX4[orig],Hu[1],Tb/+* males were crossed to *FM6/C(1)DX, y[*]f[1]* (Bloomington# 784) females.
**F_2_**: *w+ B+ non-CyO, non-TM6B* males were mated to *FM7a* (Bloomington#785) females.
**F_3_ & later:** All strains showing a white eye phenotype are discarded as insertions on autosomes. This is the easiest to discern in F_4_ *non-FM7a* males.

### Insertion site cloning

We applied an inverse PCR (iPCR) approach (Kockel et al., 2019), to molecularly clone the insertion sites of Stan-X SX4 P-elements. DNA restriction enzymes used: Sau3AI (NEB R0169) and HpaII, (NEB R0171). Ligase used: T4 DNA Ligase (NEB M0202). 5’ end cloning: Inverse PCR primer “Plac1” CAC CCA AGG CTC TGC TCC CAC AAT and “Plac4” ACT GTG CGT TAG GTC CTG TTC ATT GTT. Sequencing primer 5’ end: “SP1” ACA CAA CCT TTC CTC TCA ACA. *3’* end cloning: Primer pair “Anna” CGC AAA GCT AAT TCA TGC AGC and “SP1Berta” ACA CAA CCT TTC CTC TCA ACA AAA GTC GAT GTC TCT TGC CGA. Sequencing primer 3’ end: “SP1” ACA CAA CCT TTC CTC TCA ACA. For insertions where the sequence of one end only could be determined by inverse PCR, we pursued a gene-specific PCR approach (Ballinger and Benzer 1989) using P-element and gene-specific primers. 5’ end specific P-element primer “Chris”: GCA CAC AAC CTT TCC TCT CAA C, sequencing primer 5’ end: “Sp1”. *3’* end specific P-element primer “Dove”: CCA CGG ACA TGC TAA GGG TTA A, sequencing primer *3’* end: “Dove”. Sequence of gene-specific primers are available upon request.

### Generation of Sequence Logos and position frequency matrices (PFMs)

The construction of the SX4 sequence logo was executed as described in (Kockel et al., 2019 and Crooks et al., 2004) using http://weblogo.threeplusone.com/. The input sequence motif data is listed in Suppl. Table 1. The 8bp genomic insertion site sequence is co-directional to the P-element’s direction of insertion (Kockel et al., 2018, Linheiro and Bergman, 2008). If P-elements are inserted 5’->3’, the strand of insertion was named + (plus), and unprocessed genomic scaffold sequences as present in FlyBase were used to extract the insertion site sequences. If P-elements are inserted 3’->5’, the strand of insertion is termed – (minus), and the reverse complement of the genomic scaffold sequences were used to extract these insertion site sequences.

### Genome Sequencing

Library construction for genomic sequencing of the *w[1118], Stan-X[SX4]; iso#32[II], iso#32[III]* index line was performed separately for males and females, in two replicates each, using standard Illumina protocols. Kits used: Illumina NGS Kit Illumina® DNA Prep, (M) Tagmentation (24 Samples, IPB), #20060060, and Nextera™ DNA CD Indexes (24 Indexes, 24 Samples) #20018707. Starting material was 500ng genomic DNA isolated using the Quiagen DNeasy Blood & Tissue Kit (#69504) following the instruction for insect DNA isolation. Samples were tagmented, purified, and amplified for 5 cycles using the following Nextera DNA index adapters: Male replicate 1: H503 (i5), H710 (i7). Male replicate 2: H503 (i5) and H705 (i7). Female replicate 1: H503 (i5), H705 (i7). Female replicate 2: H505 (i5) and H705 (i7). PCR fragments were purified using Sample Purification Beads (Agencourt AMPure XP #A63880), eluted into 32ul Buffer EB (Quiagen #19086) and submitted to GeneWiz and sequenced on a Illumina HiSeq using 2×150 bp sequencing, single index. The genome sequence data of *w[1118], SX4; iso#32[II];iso#32[III]* is available on SRA https://www.ncbi.nlm.nih.gov/sra/PRJNA912892, or accession number PRJNA912892.

### Genome Sequence Data Processing and Analysis

Analysis of our whole-genome sequencing data was performed using BWA, SAMtools, and freebayes. Details of the pipeline, along with specific parameters used, are provided in the StanX_tools repository (https://github.com/sanath-2024/StanX_tools).

To use our short-read dataset to find novel, non-reference transposons **(Figure 6, Suppl. Table 2),** we deploy a similar a strategy as Linheiro et. al, 2012. We used BWA to find reads that align to both, a canonical transposon sequence as well as the FlyBase reference genome. These “split reads” were processed and sorted into groups based on alignment location and orientation. Details are provided in the StanX_tools repository (https://github.com/sanath-2024/StanX_tools). Our TE mapper represents a from the ground up multithreaded reimplementation in the Rust language, focusing on performance and simplicity.

For reproduction and verification, the sequence data is deposited on SRA (BioProject accession number PRJNA91289), and a complete build pipeline is accessible (https://github.com/sanath-2024/stan_x_paper_prep).

### Analysis of Fly Cell Atlas IPC and CC cell data

IPC and CC cell nuclei isolation from males and females was conducted in the framework of the Fly Cell Atlas (FCA, Li et al., 2022, https://www.ebi.ac.uk/biostudies/files/E-MTAB-10628/E-MTAB-10628.sdrf.txt). FASTQ sequencing files were aligned to BDGP6 version of the fly whole genome using HISAT2 (Kim et al., 2019). Single cell nuclei RNA-seq libraries representing IPCs and CC cells were filtered based on dilp2, dilp3, dilp5 and akh expression, respectively. The location of natural transposons (nTEs) and gene locations in the BDGP6 genome were taken from FlyBase. featureCounts (Yang et al., 2019), was used to assign aligned reads to transposons or genes and to obtain a count matrix for each library. When quantifying counts for nTEs, multi-mapped reads were assigned their full value to each alignment, which gives a theoretical upper bound for how much transcript could exist for a single nTE. When quantifying counts for classes of nTEs or for all TE expression (**Suppl. Fig 2C**), multi-mapped reads were assigned a value of 1/x to each alignment, where x is the number of alignments, which estimates the total amount of reads associated with the class of TE or the total number of reads coming from TEs. Count matrices were used as input to Seurat (Hao et al., 2021). Seurat VlnPlot function was used to plot unnormalized counts for gene expression (**Suppl. Fig2**).

### Training of Stan-X teachers at the Discover Now Teacher Academy

For incoming Stan-X teachers, the Stan-X Biology Course covering hybrid dysgenesis (“Module 1”), insertion site sequencing by iPCR (“Module 2”), and expression analysis in third instar larvae (“Module 3”) was offered as a two-week training course consisting of a one-week online (~3 hrs/day) session, followed by a one week session of in-person training (8 hrs/day) at the Lawrenceville School, NJ, or Stanford University School of Medicine. The two-week class was offered each year in the summer or winter. The course was staffed by instructors from participating high schools and Stanford University School of Medicine. Application deadlines and other information are detailed online at https://www.stan-x.org/.

### High School Coursework

All three Stan-X Biology Course modules are taught at Phillips Exeter Academy, NH; Commack High School, Dalton School, and Chapin School, in NY; Pritzker College Prep, and Latin School of Chicago, both in Chicago, IL; The Lawrenceville School, NJ; Lowell High School, San Francisco, CA; Loyola Marymount University, and Harvard-Westlake School, both in Los Angeles, CA; Albuquerque Academy in Albuquerque, NM; Haileybury, Hertford, U.K.; Westtown School, West Chester, PA; the Hotchkiss School, Lakeville, CT and Harvard University, Division of Continuing Education (ECPS). Students at individual schools are selected by individual schools for the Stan-X course by teachers at each respective school.

Secondary school students spent 9-10 weeks executing the hybrid dysgenesis crosses, mapping and balancing their novel SX4 strains. This was followed by 2-3 weeks for molecular determination of the SX4 insertion site, using inverse PCR and DNA sequencing using spin column-based genomic DNA preparation. The last weeks of classes are reserved for crosses with reporter strains (w; *LexAop2-CD8::GFP),* allowing for training in L3 larval dissection and epifluorescent microscopy to describe tissue specific expression patterns of novel SX4 enhancer traps.

Based on performance and recommendation of Stan-X teachers, one to three students were invited to continue studies at Stanford University School of Medicine during summer internships lasting from 2-6 weeks. These studies included further molecular mapping of transposon insertion sites, and verification of tissue patterns of enhancer trap expression. Students returning in the fall term helped instructors to run the subsequent iteration of the Stan-X class, and also pursue independent projects.

## Data and reagent availability

All Stan-X SX4 derivatives and associated data are available at the Bloomington stock center. All molecular and image data are additionally available at http://Stan-X.org. Course manuals, scaffolding problem sets and sample course daily and weekly schedules are available on request.

## Acknowledgments

We thank the Bloomington Drosophila Stock Center (NIH P40OD018537) and the TRiP at Harvard Medical School (NIH/NIGMS R01-GM084947) for transgenic fly stocks used in this study. We thank Flybase (NHGRI P41HG000739) for their contributions to the *Drosophila* research community. We thank members of the Kim group (Stanford) for advice and encouragement and welcoming summer term students. The genome sequence data of *w^1118^, SX4; iso#32^II^; iso#32^III^* is available on SRA https://www.ncbi.nlm.nih.gov/sra/PRJNA912892, or accession number PRJNA912892. K.R.C. was supported by Stanford Vice Provost Undergraduate Education and Bio-X awards. Work at Phillips Exeter Academy was supported by the John and Eileen Hessel Fund for Innovation in Science Education. We thank Glenn and Debbie Hutchins, and the Hutchins Family Foundation, for supporting innovative science research at the Lawrenceville School. Work at Westtown School was supported in part by the Dayton Coles Science Research Fund. Research at Albuquerque Academy was supported by the school’s annual fund. Work at Stanford was also supported by NIH awards (R01 DK107507; R01 DK108817; U01 DK123743; P30 DK116074), the Reid Family, Sadie and Kelly Skeff, H.L. Snyder Foundation and Elser Trust, Mr. Richard Hook, two anonymous donors, and the Stanford Diabetes Research Center (SDRC).

---

**Suppl. Table** 1: Registry of all SX4 enhancer trap insertions in this study characterized by their molecular coordinates, direction of insertion, and nearest gene. The naming is a composite of the enhancer trap construct used (SX4, Figure 1), a code of the line-generating school (Aq: Albuquerque Academy, NM. Ch: The Chapin School, NY. Co: Commack High School, NY. Da: The Dalton School, NY. Et: Phillips Exeter Academy, NH. Hb: Haileybury, U.K. La: Latin School, II. LMU: Loyola Marymount University, CA. Lv: The Lawrenceville School, NJ. Lw: Lowell High School, San Francisco, CA. Pr: Pritzker College Prep, Il. S: Stanford University School of Medicine, CA), and an internal line number. SX4 insertions that could be located to a clade of natural transposons, but the location of this natural transposon could not be unambiguously resolved within the genome, are listed at the bottom of the file, without nucleotide coordinates. The insertions into natural transposons that are not annotated in FlyBase are given a coordinate, but no FlyBase ID (FBti number).

**Suppl. Table 2:** DNA sequence of genomic library amplicons from *w^1118^, SX4; iso#32^II^; iso#32^III^* libraries spanning the breakpoints of natural TEs that are not represented in the FlyBase *Drosophila* genome release 6. The fragments were isolated by TE Mapper **(Methods),** and correspond to fragments indicated in **Figure 6.**

**Suppl. Data File** 1: Annotated DNA sequence of the SX4 element, with included color code included below. Annealing sites for inverse PCR primers and sequencing primers are marked. Regions causing BLAST hits (into white, *kirre,* and *hsp70)* in case of an empty iPCR vector are indicated.

**Suppl. Figure 1:** A) Implementation of the Stan-X curriculum during the school year and B) summer break, including a Stan-X program taught at the Department of Continuing Study at Harvard University, MA, and student internships at the Kim lab at Stanford University School of Medicine.

**Suppl. Figure 2: Expression analysis of natural TE expression on single nucleus RNA sequencing (snucRNAseq) data from corpora cardiaca (CC) and insulin expressing cells (IPCs).** A-B) Normalized expression of A) *akh* and B) dilp2 expression of filtered snucRNAseq libraries of males and females, respectively. C) Percentage of sequenced amplicons mapping to all natural TE per library. D-J) Sequenced amplicon counts present in snucRNAseq libraries of CC and IPCs for natural TEs tagged by SX4 insertions: D) *invader1{}757* (tagged by *SX4Lv807),* E) *opus{}1033* (tagged by *SX4Et49),* F) *juan{}1190* (tagged by *SX4Lv831),* G) *1360{}1206* (tagged by *SX4Lv816),* H) *F{}1209* (tagged by *SX4Lv811),* I) *mdg3{}1215* (tagged by *SX4Co822),* J) *invader4{}1371* (tagged by *SX4EXPS11).*

