## Supplementary figures and images for "An international scholastic network to generate LexA enhancer-trap lines for *Drosophila*"

### Supplemental Figure 1

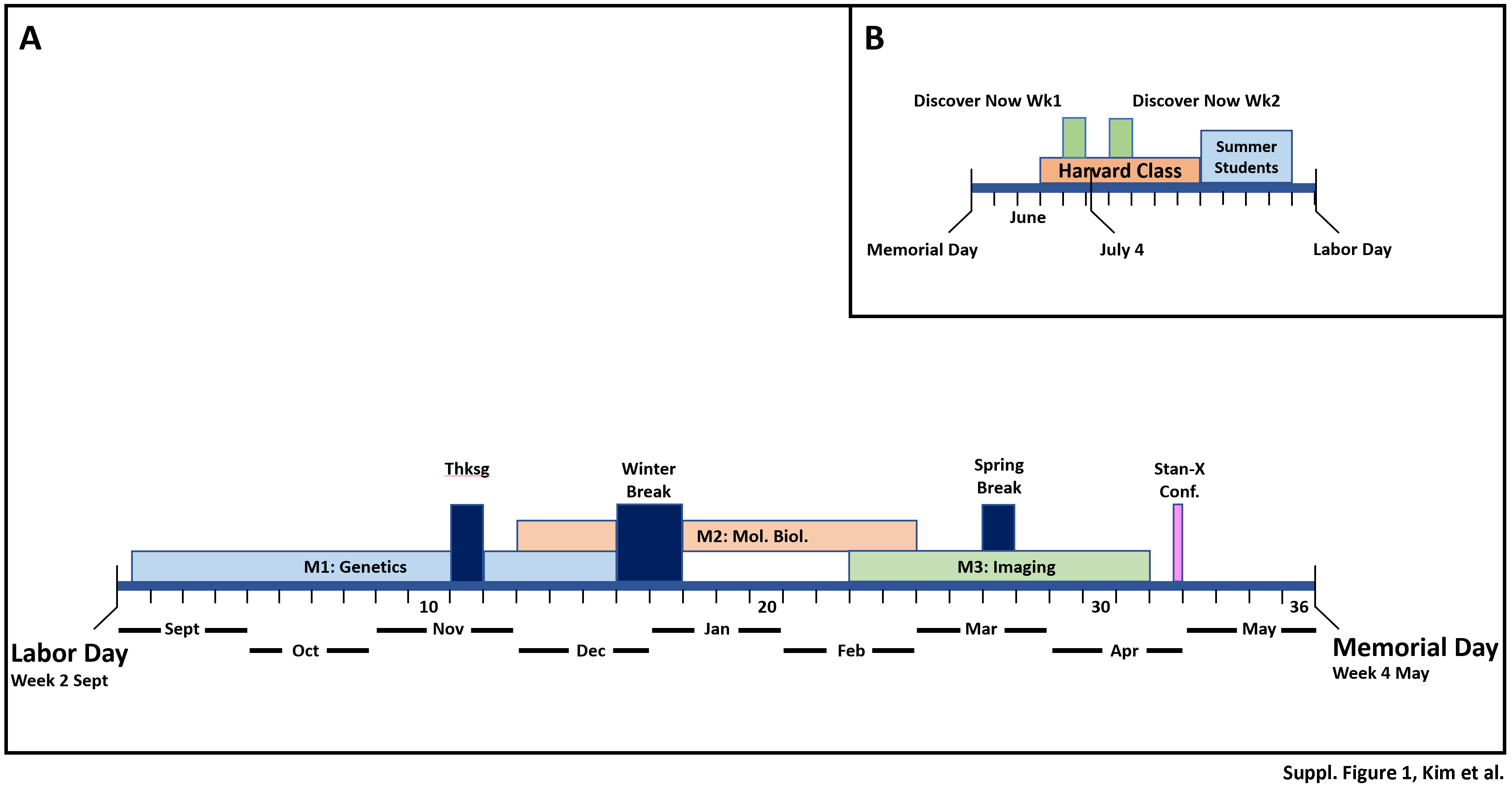

### Supplemental Figure 2

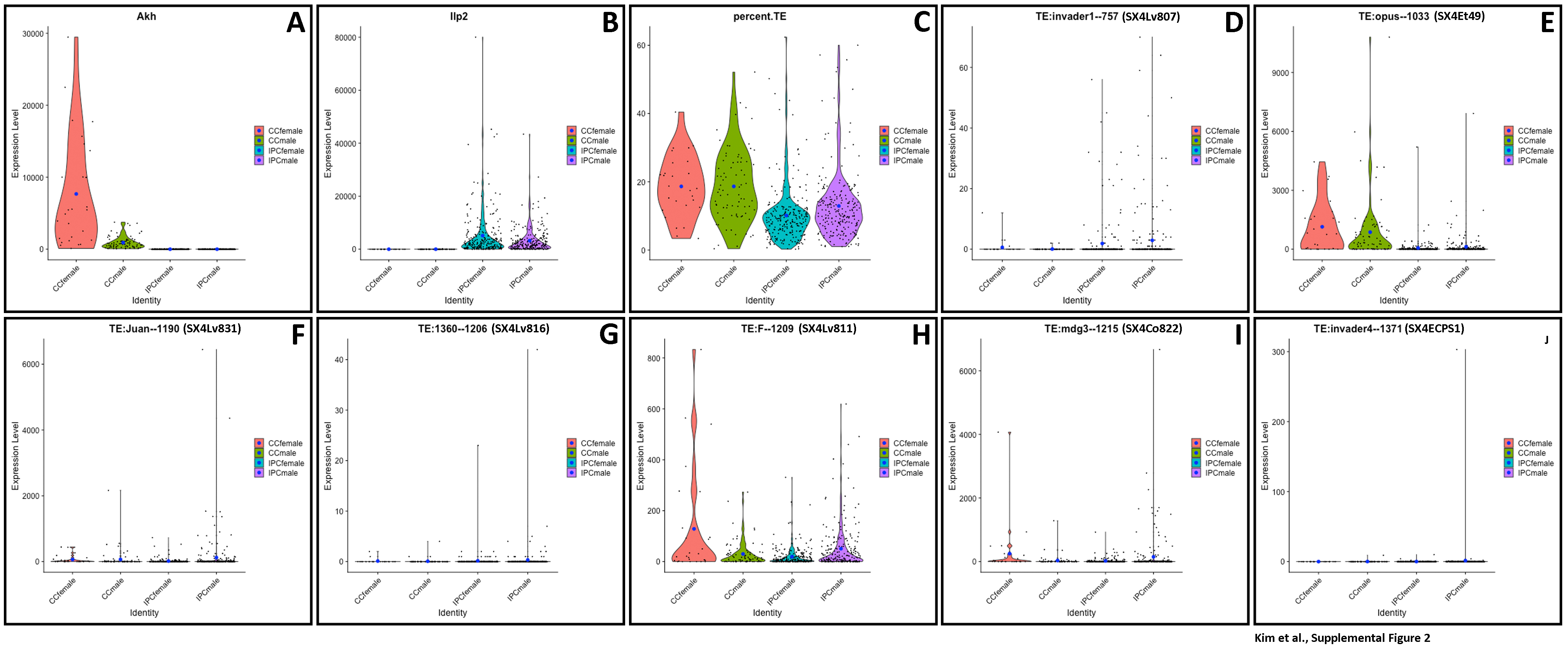
